# Human inferences about sequences: A minimal transition probability model

**DOI:** 10.1101/068346

**Authors:** Florent Meyniel, Maxime Maheu, Stanislas Dehaene

**Affiliations:** Cognitive Neuroimaging Unit, CEA DRF/I2BM, INSERM, Université Paris-Sud, Université Paris-Saclay, NeuroSpin center, Gif-sur-Yvette, France; Université Paris Descartes, Sorbonne Paris Cité, Paris, France.; Collège de France, Paris, France

**Keywords:** sequence, expectation, statistics, Bayes, inference, surprise, learning

## Abstract

The brain constantly infers the causes of the inputs it receives and uses these inferences to generate statistical expectations about future observations. Experimental evidence for these expectations and their violations include explicit reports, sequential effects on reaction times, and mismatch or surprise signals recorded in electrophysiology and functional MRI. Here, we explore the hypothesis that the brain acts as a near-optimal inference device that constantly attempts to infer the time-varying matrix of transition probabilities between the stimuli it receives, even when those stimuli are in fact fully unpredictable. This parsimonious Bayesian model, with a single free parameter, accounts for a broad range of findings on surprise signals, sequential effects and the perception of randomness. Notably, it explains the pervasive asymmetry between repetitions and alternations encountered in those studies. Our analysis suggests that a neural machinery for inferring transition probabilities lies at the core of human sequence knowledge.

Author Summary

We explore the possibility that the computation of time-varying transition probabilities may be a core building block of sequence knowledge in humans. Humans may then use these estimates to predict future observations. Expectations derived from such a model should conform to several properties. We list six such properties and we test them successfully against various experimental findings reported in distinct fields of the literature over the past century. We focus on five representative studies by other groups. Such findings include the “sequential effects” evidenced in many behavioral tasks, i.e. the pervasive fluctuations in performance induced by the recent history of observations. We also consider the “surprise-like” signals recorded in electrophysiology and even functional MRI, that are elicited by a random stream of observations. These signals are reportedly modulated in a quantitative manner by both the local and global statistics of observations. Last, we consider the notoriously biased subjective perception of randomness, i.e. whether humans think that a given sequence of observations has been generated randomly or not. Our model therefore unifies many previous findings and suggest that a neural machinery for inferring transition probabilities must lie at the core of human sequence knowledge.

## Introduction

From bird song to music, sea waves, or traffic lights, many processes in real life unfold across time and generate time series of events. Sequences of observations are therefore often underpinned by some regularity that depends on the underlying generative process. The ability to detect such sequential regularities is fundamental to adaptive behavior, and many experiments in psychology and neuroscience have assessed this ability by appealing to tasks involving sequences of events. Various effects suggestive of local sequence learning have been consistently reported, even when experimental sequences are devoid of any regularity (i.e. purely random) and restricted to only two possible items or actions. Studies of “novelty detection” for instance show that the mere exposure to a sequence of stimuli elicits reproducible “novelty” brain responses that vary quantitatively as a function of the item infrequency and divergence from previous observations [1–11]. Behaviorally, studies using two-alternative forced-choice have revealed “sequential effects”, i.e. fluctuations in performance induced by local regularities in the sequence. For instance, subjects become faster and more accurate when they encounter a pattern that repeats the same instructed response, or that alternates between two responses, and they slow down and may even err when this local pattern is discontinued [12–23]. Finally, studies asking subjects to produce random sequences or to rate the apparent “randomness” of given sequences, show a notorious underestimation of the likelihood of alternations [24–29].

Here, we propose a model that provides a principled and unifying account for those seemingly unrelated results, reported in various studies and subfields of the literature quoted above.

We adopt a Bayesian-inference approach [30–37] which relies on three pillars. The first one is that information processing in the brain relies on the computation of probabilities [30,38–43]. A second pillar is that these probabilistic computations closely approximate Bayes rule. This means that, in order to infer the hidden regularities of the inputs it receives, the brain combines the likelihood of observations given putative regularities and the prior likelihood of these regularities [44]. A third pillar is the predictive and iterative nature of Bayesian computations: once the hidden regularities of the inputs are inferred, the brain uses them to anticipate the likelihood of future observations. Comparison between expectations and actual data allows the brain to constantly update its estimates – a computational mode termed “active inference” [45,46].

To apply this general framework to sequences, one must identify the models that the brain computes when learning from a sequence. One possibility that we explore here is that there are core building blocks of sequence knowledge that the brain uses across many different domains [47]. Throughout this paper, our goal is to identify the minimal building block of sequence knowledge. By “minimal”, we mean that a simpler hypothesis would demonstrably fail to account for experimental effects such as surprise signals, sequential effects in reaction times and the biased perception of randomness.

Our proposal can be succinctly formulated: the brain constantly extracts the statistical structure of its inputs by estimating the *non-stationary transition probability matrix* between successive items. “Transition probability matrix” means that the brain attributes a specific probability to each of the possible transitions between successive items. “Non-stationary” means that the brain entertains the hypothesis that these transition probabilities may change abruptly, and constantly revises its estimates based on the most recent observations.

We formalized this proposal into a quantitative model, which we call the “local transition probability model”. As we shall see, this model predicts that expectations arising from a sequence of events should conform to several properties. We list these properties below and unpack them, one at a time, in the *Results* section.

To test whether the proposed model is general, we simulated the results of five different tasks previously published [1,2,9,20,25]. They differ in the type of observable: either reaction times [2,20], judgment of randomness [25], functional MRI signals [2] or EEG signals [1,9]. They also differ in the experimental task: either passive listening [1], two-alternative forced-choice [2,9,20] or subjective ratings [25]. Last, they also differ in the way stimuli are presented: sequential and auditory [1], sequential and visual [2,9,20] or simultaneous and visual [25]. Yet, as we shall see, all of these observations fall under the proposed local transition probability model.

We also tested the minimal character of the model, i.e. the necessity of its two main hypotheses that transition probabilities are learned and that such learning is local in time. Instead of transition probabilities, simpler models previously proposed that subjects learn the absolute frequency of items, or the frequency of alternations [9,20,23,48]. We evaluated the predictive ability of these statistics, whose relationships are illustrated in figure 1. We also tested the non-stationarity hypothesis by comparing the local transition probability model with other models that assume no change in the quantity they estimate, or a simple forgetting rule (as illustrated in Figure 2). This model comparison is not exhaustive since many proposals were formulated over the past fifty years; however, it allows to test for the necessity of our assumptions. To anticipate on the results, we found that only a learning of local transition probabilities was compatible with the large repertoire of experimental effects reported here.

**Figure 1.**
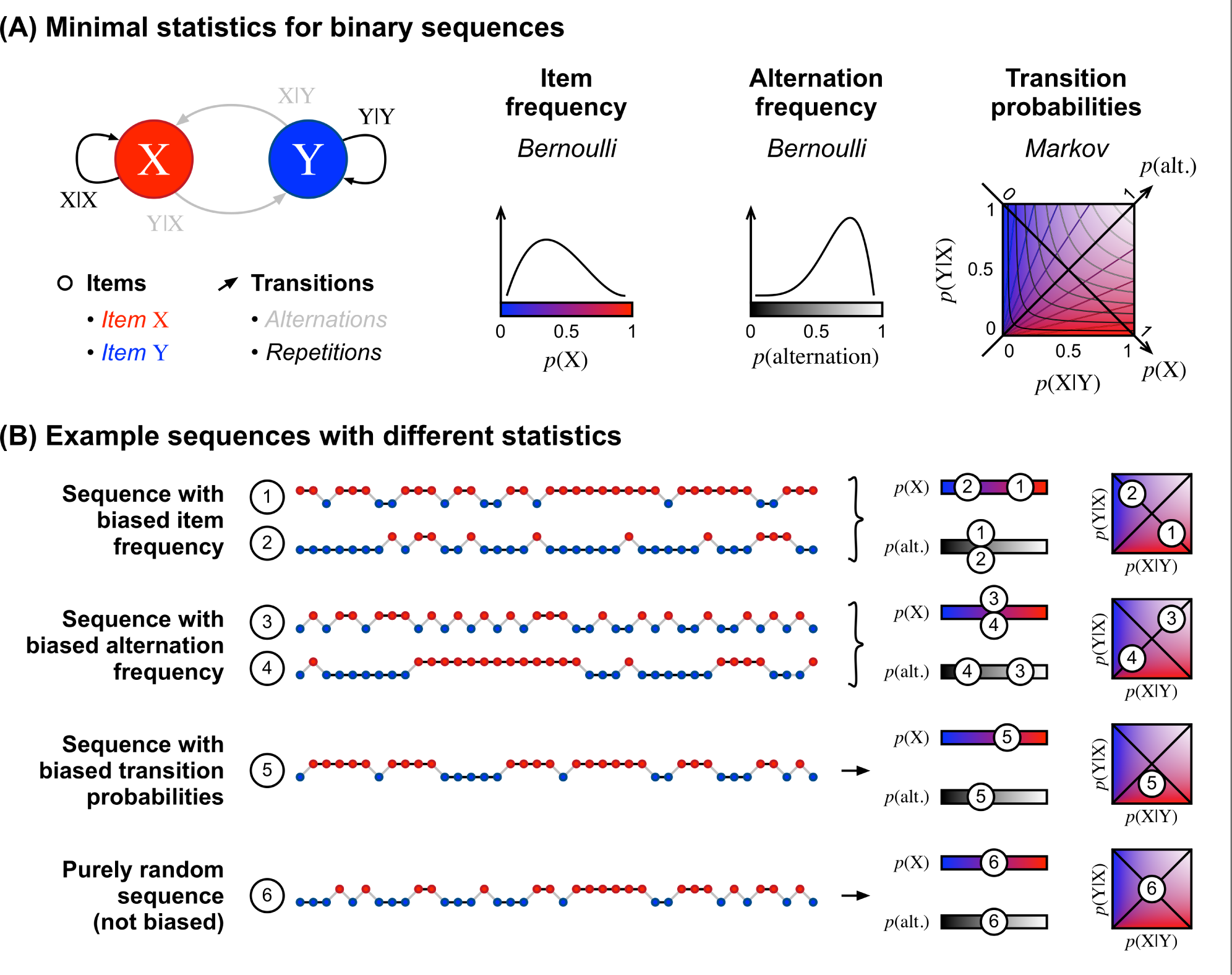
Three different hypothesis spaces. (A) Sequences can be characterized by a hierarchy of statistics. We consider here binary sequences with only two items: X and Y. The simplest statistic considers stimuli in isolation, based on the frequency of items, *p*(X) and *p*(Y). A second level considers pairs of items irrespective of their order, distinguishing pairs of identical versus different items (XX and YY vs. XY and YX). The relevant statistic is the frequency of alternations, or conversely, the frequency of repetitions: *p*(alt.) = 1 – *p*(rep.). A third level considers ordered pairs, distinguishing X_1_Y_2_ from Y_1_X_2_. The relevant statistics are the two transition probabilities between consecutive items: *p*(Y_2_|X_1_) and *p*(X_2_|Y_1_). For brevity, we generally omit the subscripts. For binary sequences, the space of transition probabilities forms a 2-dimensional matrix. In this space, the diagonals are special cases where transition probabilities coincide with the frequency of items and frequency of alternations. Out of the diagonals, there is no linear mapping between transition probabilities and the frequency of items (shown in red/blue and iso-contours) or the frequency of alternations (shown with transparency and iso-contours). Example sequences generated from distinct statistics. From top to bottom: The sequences (1) and (2) differ in their frequency of X but not in their frequency of alternations. To generate such sequences, one can select the next stimulus by flipping a biased coin. The sequences (3) and (4) differ in their frequency of alternations, but not in their frequency of X. To generate such a sequence, one can start the sequence arbitrarily with X or Y, and then decide whether to repeat the same item or not by flipping a biased coin. The sequence (5) is biased both in its frequency of alternations and its frequency of items. It cannot be generated with a single biased coin, but instead two biased coins are required, one to decide which item should follow an X and the other to decide which item should follow a Y. The sequence (6) is a purely random sequence, with no bias in either transition probabilities, and hence, no bias in item nor alternation frequencies. It can be generated by flipping a fair coin.

**Figure 2.**
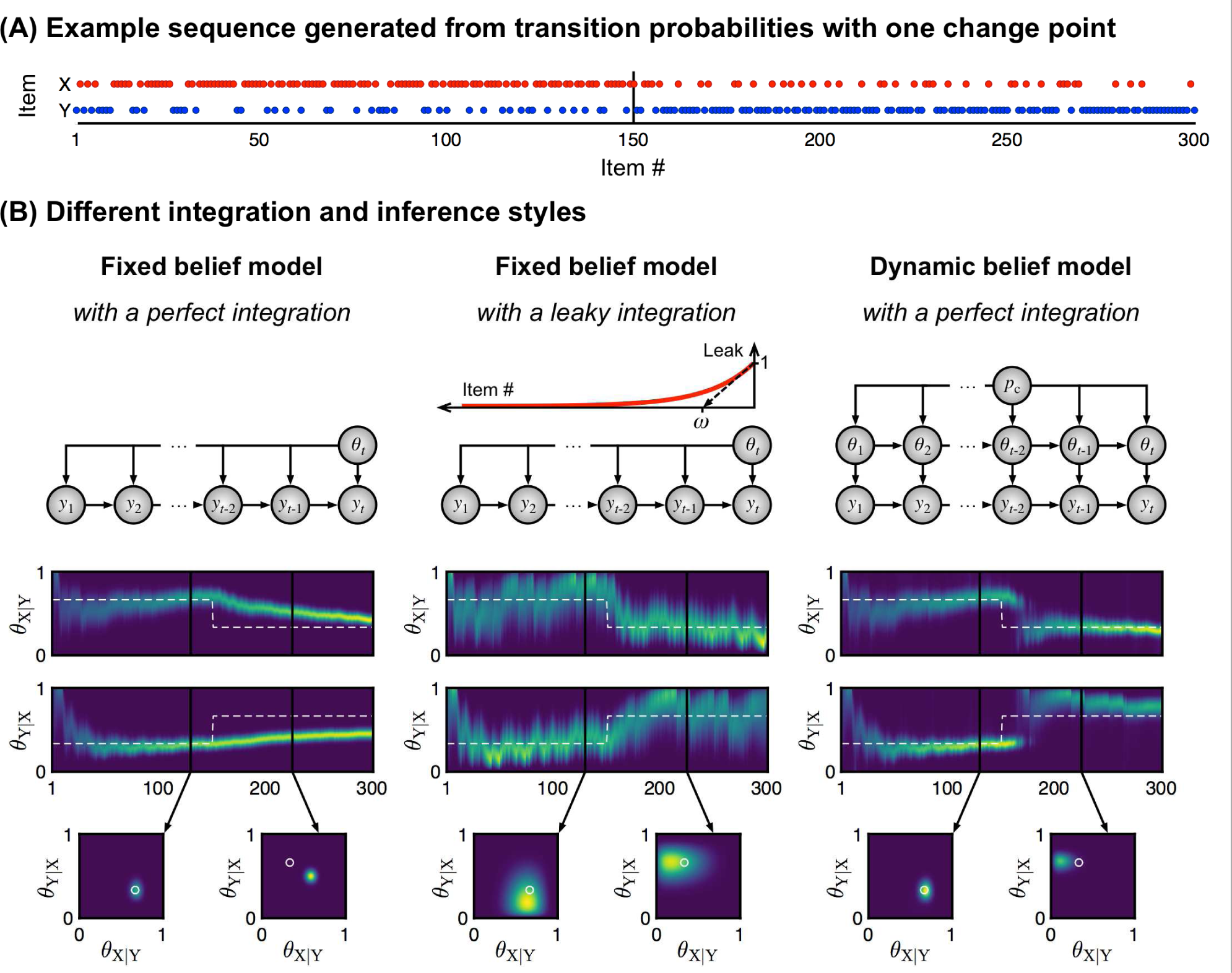
Three different inference styles. Panel A shows an example of a sequence in which the statistics change abruptly: the first half, from 1 to 150, was generated with *p*(X|Y) = 1 – *p*(Y|X) = 2/3, and the second half with *p*(X|Y) = 1 – *p*(Y|X) = 1/3. In this paper, we consider different hypotheses as to the inference algorithm used by the brain to cope with such abrupt changes (panel B). Some models assume that a single statistic generates all the observations received (“fixed belief”) while other assume volatility, i.e. that the generative statistic may change from one observation to the next with fixed probability *pc* (“dynamic belief”). Models with fixed belief may estimate the underlying statistic either by weighting all observations equally (“perfect integration”), or by considering all observations within a fixed recent window of N stimuli (“windowed integration”, not shown in the figure), or by forgetting about previous observations with an exponential decay *ω* (“leaky integration”). The heat maps show the posterior distributions of transition probabilities generating the sequence in (A) as estimated by each model. The white dash line indicates the true generative value. The insets show the estimated 2-dimensional space of transition probabilities at distinct moments in the sequence. White circles indicate the true generative values.

## Results

### Description of the model

The local transition probability model assumes that several brain circuits involved in sequence learning entertain the hypothesis that the sequence of items has been generated by a “Markovian” generative process, i.e. only the previous item y(t-1) has a predictive power onto the current item y(t). Those circuits therefore attempt to infer the “transition probability matrix” which expresses the probability of observing a given item, given the identity of the preceding one.

Further, the model is local in time in that it assumes that the transition probabilities generating the observations may change over time (some theorists call this a model with a “dynamic belief”). More precisely, it assumes that there is a fixed, non-zero probability pc that the full matrix of transition probabilities changes suddenly from one observation to the next (see Figure 2A). Therefore, at any given moment, the current and unknown generative transition probabilities must be estimated only from the observations that followed the last change. Note that the occurrence of such changes is itself unknown – the model must infer them. Bayes rule and probabilistic inference allow to solve this challenging problem optimally. Intuitively, the optimal solution discounts remote observations and adjusts the strength of this discounting process to the trial-by-trial detection of changes. The estimation of transition probabilities is therefore “local” and non-stationary.

In this paper, we contrast the local transition probability model with an alternative model which entertains a “fixed belief”, i.e. which assumes that the generative process never changes (p_c_ is exactly 0). The fixed-belief assumption greatly simplifies the estimation of transition probabilities, which boils down to counting the occurrence of each transition between any two items – but it prevents the model from adapting to the recent history of events. We also consider models which only approximate the Bayes-optimal “dynamic belief” inference. One such model is a forgetful count that discards old observations or weights recent observations more than past ones [23]. The count may be forgetful because it is limited to a fixed window of recent observations (the “windowed model”), or because it involves a leaky integration, such that previous observations are progressively forgotten. Importantly, the time scale over which forgetting occurs is fixed at a preset value and therefore cannot be adjusted to the trial-by-trial detection of changes, unlike the optimal solution.

The leaky integration and the Bayes-optimal dynamic belief are two algorithms, each with a single free parameter, that result in local estimates of statistics. Both yield similar results in the present context, we therefore refer to both as the “local transition probability model”. We reported the results for the leaky integration in the main text and the results for the Bayes-optimal dynamic belief as supplementary information.

These different inference styles of transition probabilities – fixed belief, dynamic belief, leaky integration – are depicted in Figure 2B. For comparison, we also implemented variants that resort to the same inference styles but estimate a different statistic: either the absolute frequency of items, or the frequency of alternation between successive items. It is important to note that these statistics are simple than transition probabilities, because the information about the frequency of items and the frequency of alternations is embedded in the larger space of transition probabilities (see Figure 1A). Transition probabilities also dictate the frequency of ordered pairs of items (see Supplementary Equations). Some of the models included in our comparison were proposed by others, e.g. the fixed-belief model that learns item frequencies was proposed by Mars and colleagues [4] and dynamic-belief models were also proposed by Behrens, Nassar and colleagues [49,50] for learning item frequencies, and by Yu and Cohen [23] for learning the frequency of alternations.

The local transition probability model makes several predictions:

1. *In binary sequences, expectations should reflect at least two statistics: the absolute frequency of items, and the frequency of alternations*. This is because these statistics are special cases of transition probabilities (see Figure 1A).
2. *Transition probabilities should drive expectations both globally (given the entire sequence) and locally (given the recent history of observations)*. Since the inference is local, it captures both local statistics, and their average (the global statistics).
3. *The local effects should be observed even in fully unbiased sequences where items and transitions are equiprobable*. Indeed, a local, non-stationary inference constantly captures the local deviations from equiprobability that emerge by chance even in purely random sequences.
4. *The same observations received in different orders should produce different expectations*. Since the model progressively discounts previous observations, the impact of a given observation depends on its rank within the recent past.
5. *Repeating stimuli should produce stronger expectations than the same number of alternating stimuli*. This is an emergent property of learning transition probabilities: a repeating sequence (e.g. XXXXX) provides twice as much evidence about a single transition type (X→X) than an alternating sequence (e.g. XYXYX) where the evidence is spread among two transition types (X→Y and Y→X).
6. *The perception of randomness should be biased: although, the highest degree of randomness is, by definition, achieved when there is no bias, a sequence that contains slightly more alternations than repetitions should be perceived as “more random” than a genuinely unbiased sequence*. This is because stronger expectations arise from repetition than alternation.

In the following, we characterize these predictions in greater detail in specific experimental contexts, and we test them against a variety of data sets and by comparing with simpler models.

### Effects of sequence statistics in electrophysiological data

The P300 is an event-related potential that can be easily measured on the human scalp and is sensitive to the surprise elicited by novel or unexpected stimuli. Squires et al. (1976) made a seminal contribution: they showed that, even in a random stream of items, the P300 amplitude varies strongly with the local history of recent observations. Even in purely random sequences (like fair coin flips) the amplitude of the P300 elicited by a given stimulus X increases when it is preceded by an increasing number of other stimuli Y, i.e. when it violates a “streak” of recent repetitions of Y (e.g. XXXX<u>X</u> vs. YXXX<u>X</u> vs. YYXX<u>X</u> vs. YYYX<u>X</u> vs. YYYY<u>X</u>). The P300 amplitude also increases when a stimulus violates a pattern of alternations enforced by the recent history (e.g. XYXY<u>X</u> vs. YXYX<u>X</u>). Squires et al. (1976) plotted these history effects as “trees” reflecting the entire history of recent stimuli (Figure 3A). When they varied the overall frequency of items in the sequence (from *p*(X) = 0.5 to 0.7 or 0.3), they also found that the entire tree of local effects was shifted up or down according to p(X). Altogether, their data show that the P300 amplitude reflects, in a quantitative manner, the violation of statistical expectations based on three factors: the global frequency of items, their local frequency and the local frequency of alternations. Importantly, these local effects emerged even in purely random sequences (see the middle tree in Figure 3A).

**Figure 3.**
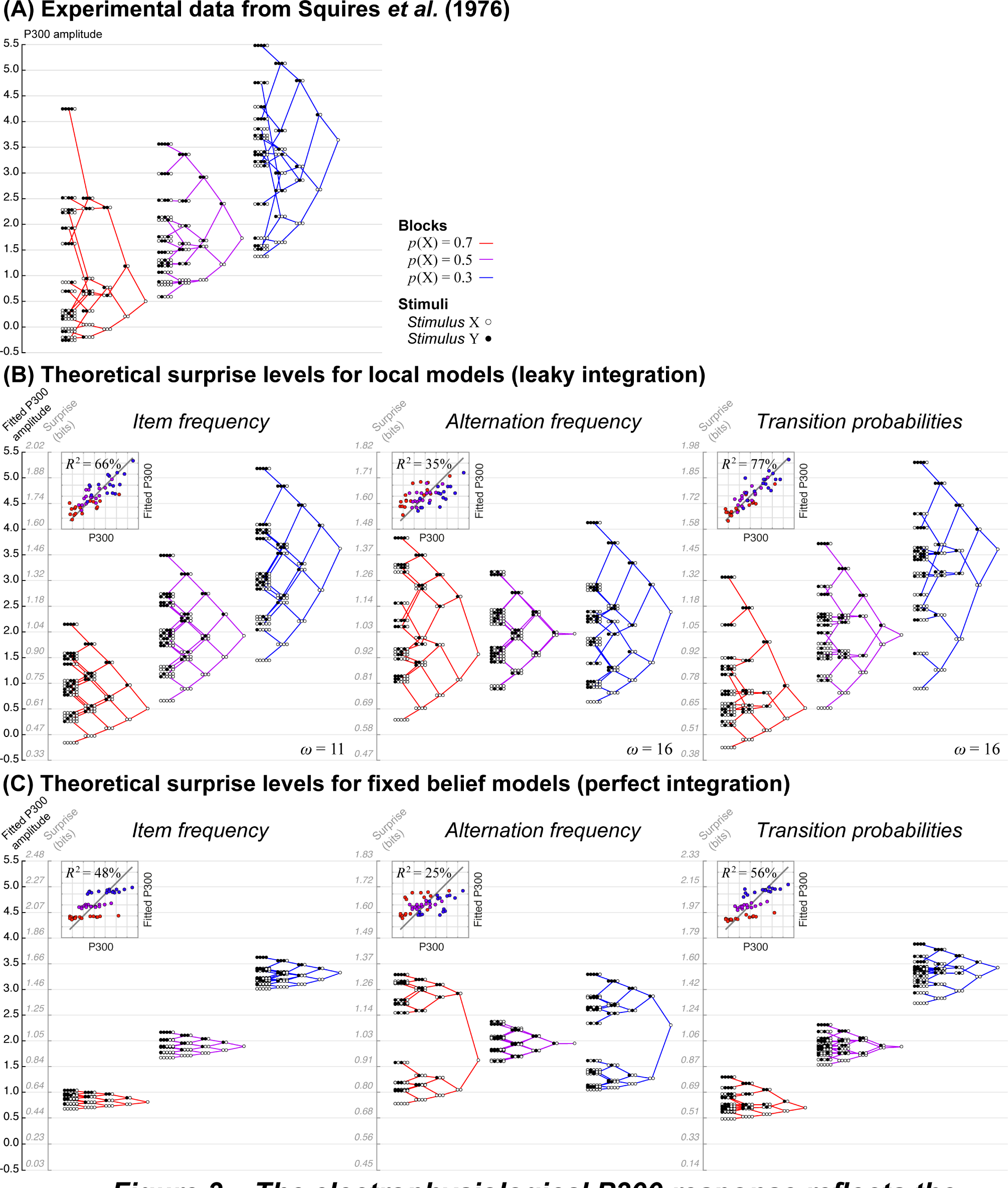
The electrophysiological P300 response reflects the tracking of statistical regularities. (A) Data redrawn from Squires et al. (1976). Subjects passively listened to binary streams of auditory stimuli (denoted X and Y). Stimuli were generated randomly with global frequency *p*(X) = 0.5 (no bias), *p*(X) = 0.7 or *p*(X) = 0.3 (biased frequencies) in separate sessions. The P300 amplitude was averaged at the end of all possible patterns of 5 stimuli at most, and plotted as a “tree” whose branches show the possible extensions for each pattern. (B-C) Average theoretical levels of surprise for all possible patterns. For each model (i.e. each set of three trees), the theoretical surprise levels were adjusted for offset and scaling to fit the data. For local models with leaky integration (B), we show the trees corresponding to the best fitting value of the leak parameter *ω*. The insets show a direct comparison between data and best-fitting theoretical surprise levels, with the regression R^2^.

These effects correspond to properties #1, #2 and #3 of the local transition probability model. Because the P300 wave seems to reflect the violation of expectations, rather than the expectations themselves, we quantified whether a given observation fulfills or deviates from expectations with the mathematical notion of surprise [51,52]. We computed theoretical levels of surprise, given the observations received (and no other information), from the local transition probability model and we found that they quantitatively reproduce the data from Squires et al. (Figure 3B).

More precisely, the local transition probability model has a single free parameter, which controls the non-stationarity of the inference. It is crucial to avoid conflating the dimensionality of the estimated quantities (which is two here, the for transition–probability matrix between two items) and the number of free parameters constraining this estimation (which is one for the local transition probability model). In the approximate model that we tested here, the only free parameter is the leak of the integration (*ω*) whose best fitting value was *ω* = 16 stimuli. This exponential decay factor means that the weight of a given observation is divided by two after a half-life of ω*ln(2)≈11 new observations. We report in SI the results for the exact inference (Bayes-optimal dynamic belief), for which the best fitting value of the a priori probability of change was *p*_c_ = 0.167.

While the assumptions of the local transition probability model seem sufficient to account for the data, we can also demonstrate that each of them is actually necessary and that they can be distinguished from one anther (see Supplementary Figure 4). Models with constant integration, i.e. without leak or a recent observation window, become increasingly insensitive to the recent history of observations as more observations are received. For such models, further details in the recent history have little impact on their expectations, as seen in the corresponding shriveled trees (see Figure 3C). Models that learn simpler statistics are also not able to fully reproduce the data. The ones that learn the frequency of alternations show little effect of the global item frequency (see the position of the roots of trees in Figure 3B). Those that learn the frequency of items capture the effect of global item frequency, but they fail to reproduce the specific arrangement of branches of the trees. For instance, in purely random sequences (when *p*(X)=0.5), such models predict that the surprise elicited by YXY<u>X</u> patterns should be like the average response (compare its position relatively to the root of the tree in Figure 3B) whereas it is not the case in the data. A model that learns transition probabilities captures the lower-than-average activity for YXY<u>X</u> patterns because it detects the repeated alternation (see Figure 3B).

We quantified the superiority of the local transition probability model using the Bayesian Information Criterion (BIC), which favors goodness-of-fit while penalizing models for their number of free parameters. The local transition probability model was better than the others: all ∆BIC > 9.46 (see Table 1). Cross-validation accuracy, another metric for model comparison, yielded the same conclusion (see Supplementary Figure 5). We also included the model proposed by Squires et al. (see Methods) that achieves a similar goodness-of-fit, but at the expanse of a higher complexity. In addition, this model is descriptive (a linear regression of several effects of interest) and not principled.

**Table 1.**
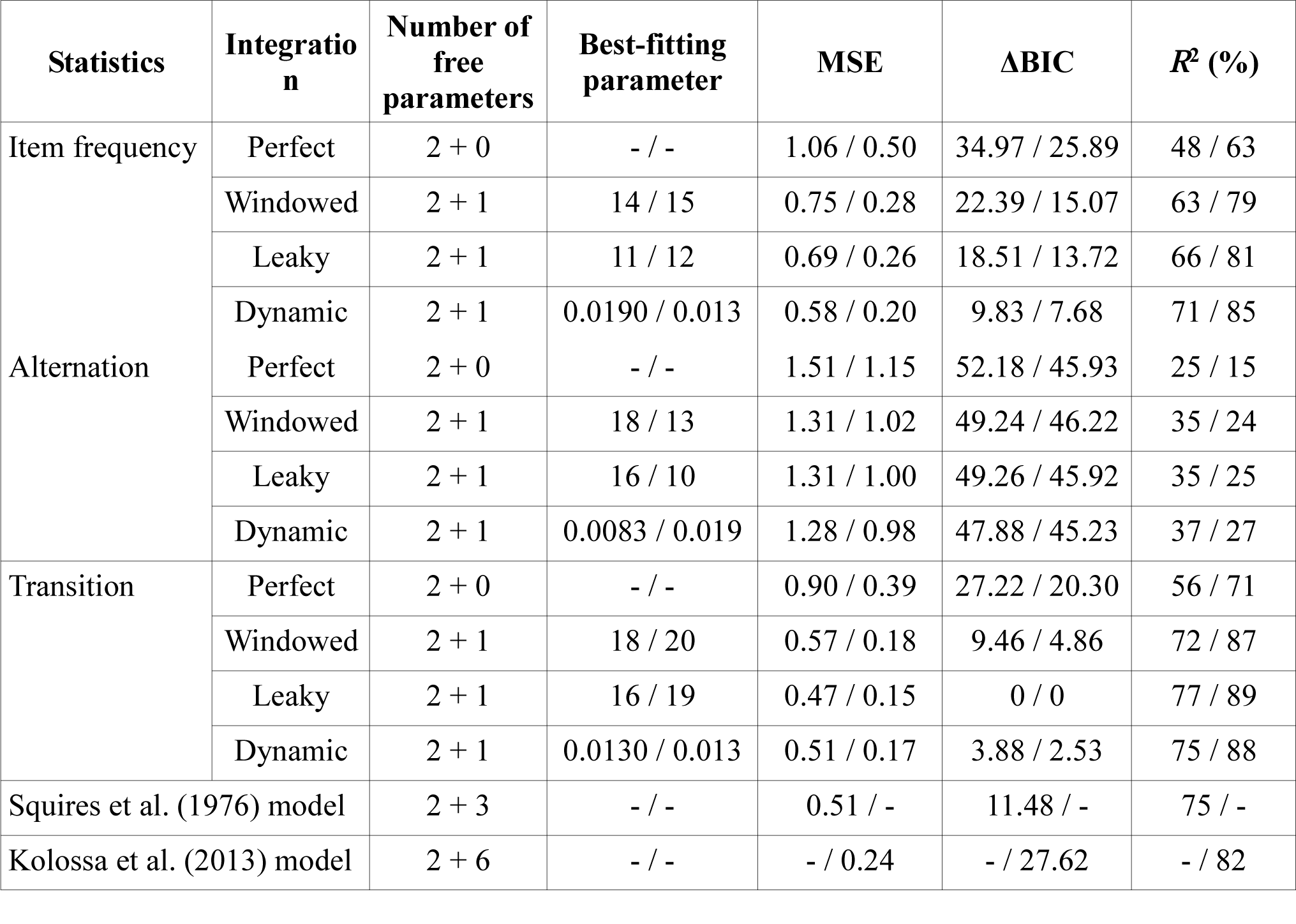
Model comparison. The table compares the fit of several models onto the dataset from Squires et al. (1976) and Kolossa et al. (2013). Values around the slash correspond to Squires / Kolossa data sets. The number of fitted parameters includes 2 linear transformations (scaling and offset) plus model-specific free parameters (from 0 to 6), see Methods for a description of the models. Best fitting parameter values are reported, excepted for the models corresponding to Squires et al 1976 and Kolossa et al. (2013) (the values are reported in the original publications) and for perfect models that have no free parameter. Bayesian Information Criterion (BIC) was computed from the mean squared error (MSE) and *n* = 48 data points (one per pattern of 5 stimuli) in Squires and *n* = 24 in Kolossa (one per pattern of 4 stimuli). BIC values are reported relatively to the best model as difference in BIC value: ∆BIC. Smaller values of MSE indicate better goodness-of-fit and smaller ∆BIC values indicate better models.

We tested the robustness of the local transition probability model with another dataset from Kolossa et al. (2013). The authors introduced noticeable differences in the original design by Squires et al.: stimuli were visual (instead of auditory) and subjects had to make a two-alternative button press for each item of the sequence (instead of listening quietly). Again, we could reproduce all qualitative and quantitative aspects of the data. Notably, we found almost the same best-fitting leak parameter (*ω* = 17 instead of 16). The BIC again favors the local transition probability model (see Table 1), even when compared to the model proposed by Kolossa et al. (see Methods).

### Sequential effects in reaction times

Reaction time tasks submit subjects to long and purely random sequences of two items. Subjects are asked to press a dedicated button for each item, and response times typically vary with the recent history of button presses. We compared subjects' reaction times to the theoretical surprise levels computed from different leaky integration models in the same experiment. Huettel et al. (2002) were interested in the effects of streaks on reaction times and brain signals recorded with fMRI. Their data show that reaction times were slower for stimuli violating a streak (see Figure 4A-B) than for those continuing it. This was true both for repeating (XXXX<u>Y</u> vs. XXXX<u>X</u>) and alternating streaks (XYXY<u>X</u> vs. XYXY<u>Y</u>), with a correlation with the streak length: the longer the streak, the larger the difference between violation and continuation. Importantly, the violation vs. continuation difference in reaction times increased more steeply with the length of repeating streaks compared to alternating streaks. This corresponds to property #5 of the local transition probability model.

**Figure 4.**
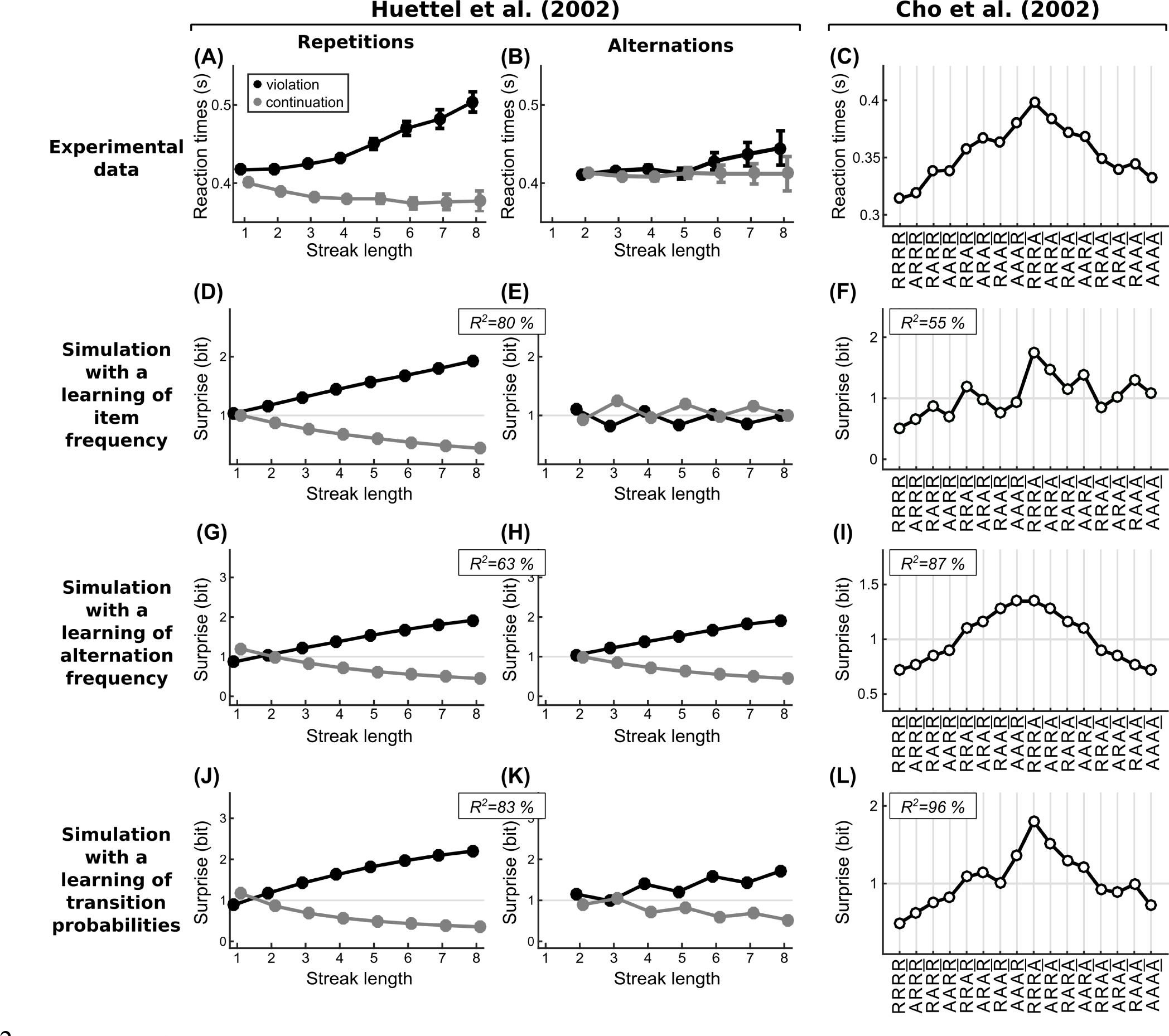
Tracking of statistical regularities and reaction times. (A-B) Experimental data redrawn from Huettel et al. (2002) [2]. Subjects were presented with a purely random stream of two items. They had to press a key corresponding to the presented item as fast as possible. Reactions times are sorted depending on whether the local sequence of items followed a local streak of repeated or alternated items, and whether the last item continued or violated the preceding pattern. For instance in XXXX<u>Y</u>, the last item violates a previous streak of four repeated items. (C) Experimental data redrawn from Cho et al. (2002) [20]. The task was similar to Huettel et al. (2002) but reaction times are now sorted based on all possible patterns of repetition (R) or alternation across the five past stimuli. For instance, the pattern AAAR denotes that the current item is a repetition of the previous item, and that the four preceding stimuli all formed alternations (e.g. XYXY<u>Y</u>). (D-L) Theoretical surprise levels estimated in purely random sequences by three different local models. These local models differ only in the statistic they estimate. Their single free parameter is the leak of integration, it was fitted to each dataset. We report the regression R^2^ for these best parameters. Note that regressions include the data for both repetitions and alternations in the case Huettel et al. Note that only a learning of transition probabilities predicts several aspects of the experimental data.

Importantly, this property is specific to a model that learns transition probabilities. A model that learns the frequency of alternations has identical expectations for repeating and for alternating streaks, because alternations and repetitions play symmetrical roles for this statistic. A model that learns the frequency of items has expectations in repeating streaks but not in alternation streaks. Indeed, as the streak length increases, the frequency of the repeated item increases but in alternating streaks, the frequency of either item remains similar. On the contrary, in a model that learns transition probabilities, expectations build up for both streak types by counting all possible transition types between successive items. In that case, an asymmetry emerges because repeating sequences offer twice the evidence about the current transition than do alternating sequences. For instance, in XXXXXXX, one may predict that the item following the last X should be another X, since six transitions X→X preceded without a single X→Y transition. In XYXYXYX, one can predict that the item following the last X should be a Y, since three transitions X→Y preceded without a single X→X transition. However, the ratio of evidence supporting the transition currently expected is stronger in the repeating sequence (6:0) compared to the alternating sequence (3:0).

One could argue that such an asymmetry is not a property of statistical learning but a simple consequence of motor constraints or motor priming. However, such a conclusion would be inconsistent with the EEG data recorded from passive subjects in Squires et al. study, in which the P300 difference between XXXX<u>Y</u> and XXXX<u>X</u> was also larger than between XYXY<u>Y</u> and XYXY<u>X</u>. In addition, Huettel et al. also recorded fMRI signals while participants performed the task in a scanner. Activity levels in several non-motor brain regions such as the insula and the inferior frontal gyrus showed the same sequential effects as the reaction times, again with a larger brain activation for violations of repetition patterns than for violations of alternation patterns.

Cho et al. (2002) were interested not only in the effect of the preceding number of repetitions and alternations on reaction times, but also in their order. To do so, they sorted reaction times based on all patterns of five consecutive stimuli (see Figure 4C). Each pattern contains four successive pairs, which can either be an alternation (denoted A) or a repetition (denoted R) of the same item. There are in total 2^4^ = 16 possible patterns of repetition and alternation. Their analysis confirmed several effects already mentioned above, such as the effect of local frequency (e.g. RRR<u>R</u> vs. RRA<u>R</u> vs. RAA<u>R</u> vs. AAA<u>R</u>).

Their data also show clear evidence for property #4 of the local transition probability model: the same observations, in a different order, produce different expectations. Consider, for instance the sequences ARR<u>R</u>, RAR<u>R</u> and RRA<u>R</u>. The local frequency of R is the same in these three patterns since they each contain a single discrepant observation (A); yet, the order of the observations matters: reaction times are slower when the discrepant A was observed more recently. In the local transition probability model, it is due to the non-stationarity of the estimation, which weights recent observations more than remote ones.

This order effect could also be reproduced by a model that learns the frequency of alternation (see Figure 4I). However, this model predicts that surprise levels should be symmetrical for alternations and repetitions. This contradicts property #5, according to which expectations build up more rapidly for alternations than repetitions. The data conform to this property: reaction times for patterns ending with a repetition are lower than those ending with an alternation (see Figure 4C), similarly to surprise levels in the local transition probability model (see Figure 4L).

Interestingly, the local transition probability model also captures additional aspects of the data that are left unexplained by a model that learns the frequency of alternations. When patterns are ordered as in Figure 4C, reaction times show gradual increases over the first eight patterns and gradual decreases for the last eight. There are also local deviations from this global trend: it is particularly salient for patterns RAA<u>R</u> and ARA<u>A</u>. The local transition probability model reproduces these local deviations. A model learning the frequency of alternations also predicts local deviations, but for other patterns (RRAR and AARA). The observed deviations are thus specific to a learning of transition probabilities. RAA<u>R</u> corresponds to XXYX<u>X</u> where the last pair XX was already observed once, whereas in ARA<u>R</u>, which corresponds to XYYX<u>X</u>, the last pair XX was not observed. Surprise is therefore lower (and not higher, as predicted by a model learning alternation frequency) for RAA<u>R</u> than ARA<u>R</u>. A similar explanation holds for ARA<u>A</u> vs. RAA<u>A</u>.

Finally, note that a model that learns the frequency of items fails to reproduce many aspects of the data (see Figure 4F) since it is completely insensitive to repetition vs. alternation effects.

We obtained the results shown in Figure 4 by fitting the leak parameter *ω* of each model. The best-fitting value for Huettel / Cho data was: *ω*=8 / *ω*=4 with a learning of stimulus frequency, *ω*=6 / *ω*=1 with alternation frequency, and *ω*=6 / *ω*=3 with transition probabilities. However, simulations using the leak value fitted to the independent dataset by Squires et al (Figure 3) led to the same qualitative conclusions. Thus, a single set of parameters may capture both data sets.

### Asymmetric perception of randomness

The asymmetry in expectation for alternation vs. repetition is probably the least trivial property of the local transition probability model (#5). This property is evidenced above in sequential effects and it entails a prediction in another domain: judgments of randomness should also be asymmetric. This prediction is confirmed: the human perception of randomness is notoriously asymmetric, as shown in particular by Falk & Konold (1997) (see Figure 5A). Sequences with probabilities of alternations *p*(alt.) that are slightly larger than 0.5 are perceived as more random than they truly are. This is an illusion of randomness: in actuality, the least predictable sequence is when *p*(alt.) = 0.5, i.e. when the next item has the same probability of being identical or different from the previous one.

**Figure 5.**
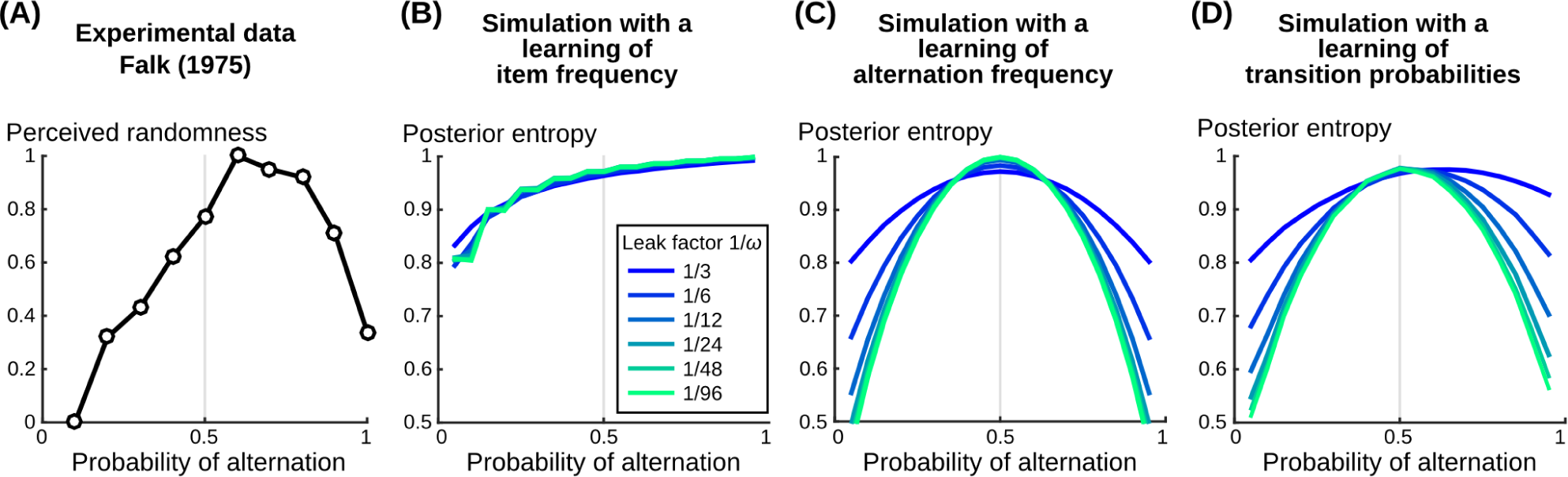
Asymmetric perception of randomness. (A) Data redrawn from Falk (1975) and reported in [25]. Subjects were presented with various binary sequences of 21 stimuli. They were asked to rate the apparent randomness of each sequence. The range of perceived randomness was normalized between 0 and 1. Ratings were sorted based on the alternation frequency in the sequences. (B-D) Theoretical levels of entropy estimated by distinct local models. The entropy characterizes the unpredictability of a sequence. For each model, we generated random sequences differing in their alternation frequencies. For each sequence and model, we computed the estimated probability of the next stimulus of the sequence, given the preceding stimuli. We then converted these predictions into entropy levels and plotted the average for different values of the leak parameter of the model. Note that only a learning of transition probabilities predicts a slight asymmetry of perceived randomness.

This bias in the perception of randomness is actually rational from the viewpoint of the local transition probability model. In order to quantify the perceived randomness of a sequence in the local transition probability model, we estimated the unpredictability of the next outcome. This unpredictability is formalized mathematically by the notion of entropy. The resulting estimated entropy level was maximal for sequences with *p*(alt.) larger than 0.5 (see Figure 5D). This bias was all the more pronounced that fewer stimuli were taken into account in the estimation: a model with a stronger leak results in a larger bias. This aspect is specific to the local transition probability model. In contrast, a model that learns the frequency of alternation shows no bias because alternations and repetitions play symmetrical roles for such a model (see Figure 5C). On the other hand, a model that learns the frequency of items shows an extreme bias: the maximal entropy level is reached for *p*(alt.) = 1 (see Figure 5B). This is because when stimuli alternate, their observed frequencies are identical, closest to chance level (50%) from the point of view of an observer that focuses solely on item frequency.

To understand how the asymmetry emerges, one should note that, in the local transition probability model, expectations arise from both repeating transitions (XX and YY) and alternating transitions (XY and YX). High expectations arise when one transition type is much more frequent than the other. The estimated entropy therefore decreases when *p*(alt.) approaches 1, where alternating transitions dominate, and when *p*(alt.) approaches 0, where repeating transitions dominate. However, remember that stronger expectations arise from repetitions than alternations in the local transition probability model (property #5). Therefore, expectations are not symmetric with respect to *p*(alt.), but higher for p(alt)<0.5 than p(alt)>0.5, so that the ensuing estimated entropy peaks at a value of *p*(alt.) that is slightly higher than 0.5.

This asymmetry is also dampened, without being abolished, when the leaky integration parameter of the local transition probability model is weaker. Indeed, experimental evidence confirms that the difference in expectations arising from repeating and alternating transitions is more pronounced for shorter sequences (see the results from Huettel et al, Figure 4A-B).

## Discussion

We showed that learning non-stationary transition probabilities entails six properties. First, expectations derived from such a learning show effects of both the frequencies of items and their alternations because these statistics are specific aspects of transition probabilities (#1). Second, these effects emerge both globally and locally in the learning process because the inference is non-stationary (#2). Third, this non-stationarity also entails that local effects emerge even in purely random sequences (#3). Fourth, it depends on the exact order of observations within the local history (#4). Fifth, since the space of transition probabilities is more general than the frequencies of items and their alternations, the local transition probability model makes a non-trivial prediction, unaccounted for by simpler statistics: expectations build up more strongly from repetitions than from alternations (#5). Sixth, this asymmetry translates into a subjective illusion of randomness which is biased toward alternations (#6). We identified many signatures of expectations and their violation in human behavior (such as reactions times) and brain signals (measured by electrophysiology and fMRI) which conformed both qualitatively and quantitatively to these predictions. We therefore conclude that transition probabilities constitute a core building block of sequence knowledge in the brain, which applies to a variety of sensory modalities and experimental situations.

Early studies [14,16] proposed that the information provided by stimuli modulates reaction times within sequences [12]. According to the information theory framework, an observation is informative inasmuch it cannot be predicted [51]. In line with this information-theoretic approach, the local transition probability model quantifies the extent to which an observation deviates from the preceding ones. The central role of expectations in cognitive processes has already been put forward by the predictive coding [7,53] and the active inference [46,54] frameworks, and applied, for instance, to motor control [55,56] or reinforcement learning [57].

However, some have claimed that sequential effects in reaction times arise from low-level processes such as motor adaptation. For instance, Bertelson wrote in 1961 “one must thus admit that the shorter reaction times [for repetitions] cannot depend on something which must be learnt about the series of signals – unless one assumes that this learning is fast enough to be completed and give already its full effect on performance in the first 50 responses” [13]. In contrast, the local transition probability model shows that, with optimal statistical learning, sequence effects can arise from a very local integration: our fit of Squires et al. (1976) data suggests a leak factor ω of 16 stimuli, meaning that the weight of a given observation is reduced by half after 16*ln(2)≈11 observations. In addition, facilitation of reaction times is observed for both streaks of repetitions and streaks of alternations, which speaks against a pure motor interpretation [17,18]. Moreover, similar sequential effects are also observed in electrophysiological and fMRI measures of brain activity in the absence of any motor task. Therefore, although motor constraints may also contribute to reaction time fluctuations, a parsimonious and general explanation for sequential effects is that they arise from learned statistical expectations.

The non-stationary integration also explains why both local and global effects emerge and why local effects persist in the long run even within purely random sequences [20,23]. From the brain's perspective, the constant attempt to learn the non-stationary structure of the world could be a fundamental consequence of a general belief that the world can change at unpredictable times, as already suggested by others [23]. Many studies indeed show that the brain can perform non-stationary estimation and thereby efficiently adapt to changes in the environment [49,50,58–60]. Technically, the belief in a changing world can be captured in two different ways: either by the *a priori* likelihood of a sudden change (a.k.a. volatility) *p*_*c*_ in the exact dynamic belief model, or by the leaky integration factor *ω* in the approximate model. The present data do not suffice to separate those two possibilities. This is because the latter (leaky integration) is such a good approximation of the former that both are difficult to disentangle in practice. Leaky integration is a popular model in neuroscience because it seems easy to implement in biological systems [23,58,61,62]. However, the dynamic belief model may not be less plausible given that neuronal populations have been proposed to represent and compute with full probability distributions [33,41]. Furthermore, only the full Bayesian model recovers an explicit probabilistic representation of change likelihood and change times. Several recent experimental studies suggest that the brain is indeed capable of estimating a hierarchical model of the environment, and that human subjects can explicitly report sudden changes in sequence statistics [60,63].

Our results suggest that, during sequence learning, the brain considers a hypothesis space that is more general than previously thought. We found that sequential effects in binary sequences are better explained by a learning of transition probabilities (a 2-dimensional hypothesis space) than of the absolute item frequencies or the frequency of their alternations (which are one-dimensional spaces). Importantly, all of these models have the same number of free parameters, so that the local transition probability model is more general without being more complex or less constrained. The critical difference lies in the content of what is learned (e.g. item frequencies vs. transition probabilities). More is learned in the latter case (a 2D space is larger than a 1D space) without resorting to any additional free parameter. The value of the learned statistic is not a free parameter, it is instead dictated by the sequence of observations and the assumptions of the model. In general, a Bayesian learner may consider a vast hypothesis space (see the many grammars used by Kemp and Tenebaum [64]) and yet, as a model that attempts to capture human behavior, it may possess very few or even zero adjustable parameters.

An alternative to the full 2D transition-probability model would be to combine two learning processes: one for the frequency of items and one for the frequency of alternations. However, such a model introduces a new free parameter compared to the local transition probability model: the relative weight between the predictions based on the frequency of items and the predictions based on the frequency of alternations. In addition, the distinction between learning transition probabilities vs. the frequency of items and their alternations is not a simple change of viewpoint: the correspondence between the two is extremely non-linear as shown in Figure 1A. Learning the frequency of items and the frequency of alternations is therefore not only less parsimonious than learning transition probabilities, it is also genuinely different.

The difference between learning transition probabilities vs. the frequency of items and their alternation may have been overlooked in the past. However, the distinction is important since these learning strategies make distinct predictions about the asymmetry of expectations arising from repetitions and alternations. This asymmetry is a classical aspect of data, in particular response times [13]. In previous models, this asymmetry was simply assumed and incorporated as a prior [23,25]. We show here, to our knowledge for the first time, how this asymmetry follows naturally from first principles (Bayes’ rule) in the local transition probability model. Moreover, our account is also unifying since it addresses not only sequential effects but also judgments of randomness.

We claim that the learning of transition probabilities is a core and general building block of sequence knowledge because we found supportive evidence in five representative datasets. There is also additional evidence from other fields. For instance, word segmentation in language relies on transition probabilities between syllables [65]. Moreover, neurons in the monkey inferior temporal cortex reduce their firing in direct proportion to the learned transition probabilities [66,67]. Ramachandran et al (2016), in particular, present single-cell recordings suggesting that the expectation about the next item does not depend on its absolute frequency or the absolute frequency of the pair it forms with the previous item, but instead on the conditional probabilities of items learned with a covariance-based rule. Additional sources of evidence that human subjects learn transition probabilities is provided by studies of “repetition suppression” [68,69], choices in decision-making problems [70] and explicit reports of learned transition probabilities [60]. The study by Bornstein and Daw [70] in particular shows that humans can learn transition probabilities among 4 items. Our local transition probability model naturally extends from the binary case to a larger number of categories, a situation that is pervasive in every-day life. Learning only the frequency of items and of repetitions becomes gradually inadequate when the number of items increases, because most environmental regularities are captured by various item-specific transition probabilities rather than absolute frequencies. For instance, the probability of imminent rain is typically high after a thunderstorm, but very low during a sunny day, and intermediate in case of strong wind. The learning of transition probabilities may even operate without awareness [71–73].

Our claim that a learning of transition probabilities accounts for a variety of experimental effects does not rule out the possibility that the brain also computes simpler statistics. Many studies report effects of item frequencies or alternation frequency. Electrophysiology in particular shows that these effects unfold across time and across brain circuits, as reflected in signals such as the mismatch negativity and the P300 [6,7,11,19]. In particular, Strauss et al. (2015, experiment 2) identified two distinct time windows in magneto-encephalographic recordings, during which the absolute frequency of items and the frequency of alternations, respectively, affected the human brain responses to simple sounds. However, in most studies, it is not clear whether such effects are particular cases of a general learning of transition probabilities or whether they are genuinely limited to item frequency or alternation frequency. Both hypotheses are indistinguishable in most studies because of their experimental design. Therefore, it is not clear for the moment whether different brain circuits are tuned to these different statistics and compute them in parallel [48], or whether most brain regions are equipped for the computation of transition probabilities. By contrast, it seems that more sophisticated building blocks of sequence knowledge, such as ordinal knowledge, chunking, algebraic patterns and tree structures are operated by specific brain circuits [47]. Future work should aim to incorporate these additional levels of representation to the local transition probability model, which we propose here as a minimal building block, likely to be duplicated in many brain regions and shared by humans and other animals alike.

## Methods

We provide the MATLAB code for our ideal observer models to reproduce our simulations and figures: https://github.com/florentmeyniel/MinimalTransitionProbsModel. In particular the “trees” corresponding to Figure 3 and simulated with the dynamic belief model are not shown in the article, but they can be easily generated with our code.

### Formal description of the models

The models are “ideal observers”: they use Bayes rule to estimate the posterior distribution of the statistic they estimate, θ_t_, based on a prior on this statistic and the likelihood provided by previous observations, *y*_*1:t*_ (here, a sequence of Xs and Ys). Subscripts denote the observation number within a sequence.

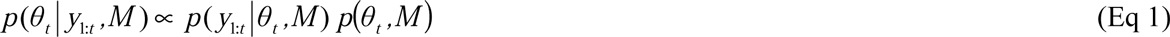

Different ideal observer models (M) estimate different statistics. The parameter θ can be the frequency of items, the frequency of alternations, or transition probabilities between items. The estimation of θ depends on the assumption of the ideal observer model: it can either consider that θ is fixed and must generate all the observations (“fixed belief models”) or that θ may change from one observation to the next (“dynamic belief models”). For all models, we use a prior distribution that is non-informative: all possible values of θ are considered with equal probability.

Note that a model estimating the frequency of alternations is equivalent to a model estimating the frequency of items after recoding of the stimuli as repetitions or alternations. Therefore, we only present below the derivation for the item frequency and transitions probabilities, in the case of both fixed belief and dynamic belief models.

#### Fixed belief models

For fixed belief, θ should not depend on the observation number. Therefore, the likelihood function can be decomposed as follows using the chain rule:
 
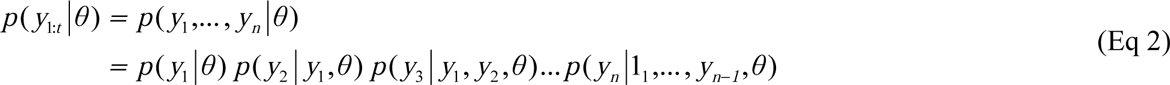

For models that estimate the frequency of items, the likelihood of a given observation depends only on the estimated frequency:

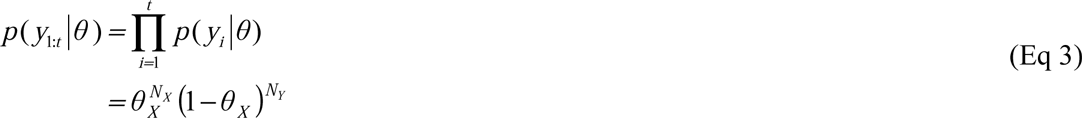

Where *θ*_X_ is the frequency of X, *N*_X_ and *N*_Y_ the numbers of X and Y in the sequence *y*_1:t_. This likelihood is a Beta distribution whose parameters are *N*_X_ + 1 and *N*_Y_ + 1. In order to derive the posterior, the likelihood must be multiplied by the prior. Here, the prior is a non-informative, flat distribution: a Beta distribution with parameters [1, 1]. Since the product of two Beta distributions is also a Beta distribution in which the parameters are simply added, the posterior distribution has an analytical solution:

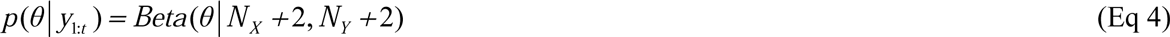

For models that estimate transition probabilities between consecutive stimuli, the likelihood of a given observation depends only on the estimated transition probabilities and the previous stimulus:

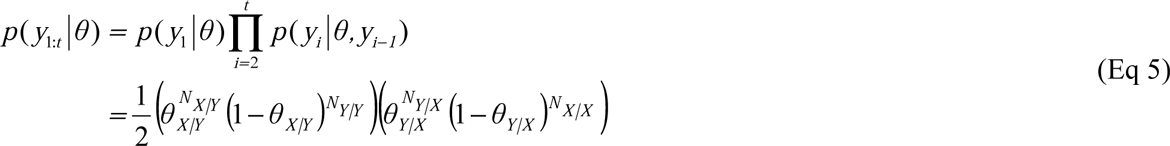

Where θ denotes a vector of two transition probabilities θ = [θ_X|Y_, θ_Y|X_], and *N*_X|Y_ denotes the number of YX pairs in the sequence *y*_1:t_. For simplicity, the first observation can be considered as arbitrary, so that *p*(*y*_1_|θ) = 1/2. Equation 5 therefore corresponds to the product of two Beta distributions, with parameters corresponding to the transition counts plus one. The product of this likelihood and a uniform prior distribution results in an analytical solution:

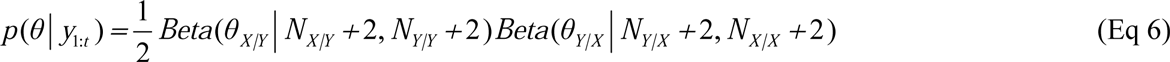

For models that estimate frequency and transition probabilities, we distinguish between different ways of counting. The leaky integration was modeled with a free parameter: an exponential decay ω on the previous observations, i.e. a weight *e*^*–k/ω*^ for the *k*-th past stimulus. The perfect integration was modeled by counting all stimuli equally. For the sake of completeness, we also included a model that counts perfectly in a window of recent observations. The length of this window is a free parameter. Note that “perfect integration” is a special case of the window size being equal or larger than the number of stimuli in the sequence.

The posterior distribution can then be turned into the likelihood of the next stimulus using Bayes rule (again):

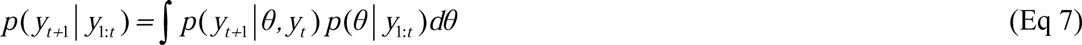

Note that for models that learn frequencies, θ is a single number (and not a vector, as for transition probabilities) and the conditional dependence on *y*_*t*_ can be ignored.

#### Dynamic belief models

In dynamic belief models, θ may change from one observation to the next with probability *p*_c_, which is the only free parameter of the model. The computation is tractable given the so-called Markov property of the generative process. If one knows θ at time *t*, then the next observation *y*_*t*+1_ is generated with θ_*t*+1_ = θ_*t*_ if no change occurred and with another value drawn from the prior distribution otherwise. Therefore, if one knows θ_*t*_, previous observations are not needed to estimate θ_*t*+1_. Casting the generative process as a Hidden Markov Model (HMM) allows to compute the joint distribution of θ and observations iteratively, starting from the prior, and updating this distribution by moving forward in the sequence of observations:

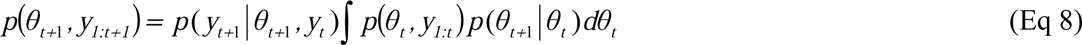

The first term in the right hand side is the likelihood of the current observation. The first term within the integral is the joint distribution from the previous iteration. The last term captures the a priori changes in θ from one observation to the next: the probability that it stays the same is 1 – *p*_*c*_ and the probability of another value is *p*_*c*_ times the prior probability of θ for that particular value. This integral can be computed numerically by discretization on a grid. The posterior probability can be obtained by normalizing the joint distribution.

The posterior distribution can then be turned into the likelihood of the next stimulus, using Bayes rule (again) and the a priori changes in θ.

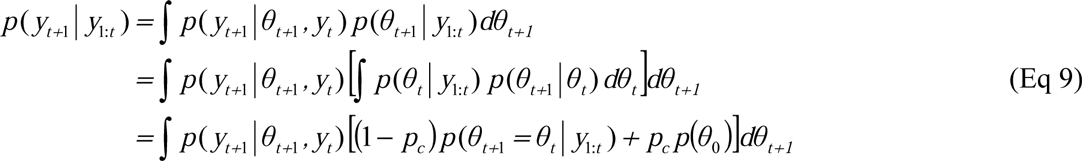

Note that for the estimation of transition probabilities, θ has two dimensions. For the estimation of frequencies, the likelihood of *y*_*t*+1_ given θ_*t*+1_ does not depend on the previous observation *y*_*t*_ so that *y*_*t*_ should be omitted on the right hand side.

### Summary of the experimental procedure in Squires et al

Squires et al. (1976) presented 7 subjects with sequences of two auditory stimuli (pure tones of 1500 Hz and 1000 Hz, denoted X and Y) during electroencephalogram (EEG) recording. In separate sessions, the sequences were generated randomly with *p*(X) equals to 0.5 (no bias) or 0.7 (biased condition). In the biased condition, *p*(Y) = 1 – *p*(X) = 0.3. Because X and Y play symmetrical roles, the authors present results for a virtual condition “*p*(X) = 0.3” which actually corresponds to analyzing the responses to item Y in the biased condition. Subjects were not told about these exact probabilities. The stimulus duration was 60 ms and the stimulus onset asynchrony was 1.3 s. Subjects were asked to count the number of X items silently and report their count after each block of 200 trials. They were presented in total and in each condition, with 800 to 1600 stimuli.

Squires et al. measured a P300 score for each stimulus. This score is a weighted combination of signals measured at central electrodes (Fz, Cz, Pz) and latencies corresponding to the N200, P300 and slow wave. The weights derive from a discriminant analysis to separate optimally the signals elicited by rare and frequent patterns (XXXX<u>Y</u> vs. XXXX<u>X</u>). The average scores are reported in each condition, for all patterns of five stimuli terminated by X.

### Summary of the experimental procedure in Kolossa et al.

Kolossa et al. collected EEG data from 16 subjects who were presented with a stream of two visual stimuli (red or blue rectangles, denoted X and Y; the mapping was counterbalanced across participants). In separate blocks, stimuli were generated with *p*(X) = 0.5 (no bias condition) or 0.7 (biased condition). A virtual condition *p*(X) = 0.3 corresponds, as in Squires et al., to the response to item Y in the biased condition. Subjects were not told about these exact probabilities. Subjects completed 12 blocks of 192 stimuli with 6 blocks in a row for each condition. The order of conditions was counterbalanced across participants. The stimulus duration was 10 ms and the stimulus onset asynchrony was 1.5 s. Subjects were asked to press a dedicated button for each item as quickly and accurately as possible.

Kolossa et al. measured the P300 amplitude at electrode Pz. The exact latency of the measurement varied across trials and participants. The authors first identified subject-specific peak latencies for the difference between rare and frequent items in the biased condition. Then, in each trial they extracted the maximum value of the signal within a window of 120 ms centered on subject-specific peaks. P300 levels are reported for each condition, for all patterns of four stimuli terminated by X.

### Fitting procedure for Squires et al. and Kolossa et al. data

We generated three sequences of 200 stimuli with probabilities *p*(X) equal to 0.3, 0.5 and 0.7. For each sequence, we computed the inference of the hidden statistics for each observer model and different values of their free parameters (if any). We computed surprise, in bit of information, for each model and each stimulus in the sequences, as – log_2_(*p*(*y*_*t*_|*y*_1:*t*–1_)), where *p*(*y*_*t*_|*y*_1:*t*–1_) is the likelihood of the actual observation. We sorted surprise levels by patterns of five stimuli terminated with X. We repeated this simulation 200 times and we averaged over simulations to reach stable results.

To compare our simulation with the data from Squires et al, we extracted their values from figure 1 in [1]. For each model, we adjusted the offset and scaling to minimize the mean squared error (MSE) between simulated and experimental data. We repeated this procedure for different values of the free parameter *ω* in fixed belief models with leaky integration, and different values of *p*c in dynamic belief models. For all models, we fitted the data only for patterns of 5 stimuli since shorter patterns are not independent from longer ones: they are weighted averages of the data obtained for longer patterns. Including shorter patterns would have inflated some aspects of the data. For instance, the effect of item frequency can be seen for all pattern lengths, including length 1, but by definition, the effect of alternations can be seen only in longer patterns. Therefore, including shorter patterns would have over-weighted the effect of global item frequency relatively to local alternations.

Our fitting procedure gives the same weight to all patterns of 5 stimuli, although rare patterns are more likely to be corrupted by noise in the experimental data. However, our results are robust to this choice and are replicated when using a weighted MSE, taking into account the expected frequency of patterns. We also checked that the grids of values used for *p*_*c*_ or ω were sufficiently dense around the global maxima, as shown in Supplementary Figure 1.

We replicated this procedure with the data from Kolossa et al., taking their values from figure 8 in [9]. The only difference was that we used patterns of length 4, as reported by Kolossa et al., instead of length 5 as Squires et al.

For comparison, we implemented the models previously proposed by Squires et al. and Kolossa et al. These models are fully described in the related articles [1,9]. In short, the model by Squires et al. is a weighted sum of three factors (the variables within brackets correspond to the notations by Squires et al.):

- the global frequency of X in the sequence (P),
- the local number of X within each pattern of 5 stimuli (M),
- an alternation score for each pattern of 5 stimuli (A).

The free parameters of this model are the relative weight of global frequency (P) vs. local frequency (M), the relative weight of global frequency (P) vs. patterns of alternations (A), and a decay factor for counting the local number of stimuli (M).

The model by Kolossa et al. is a sophistication of the model by Squires et al. It is a weighted sum of three factors, thought of as the output of digital filters computing probabilities. These filters correspond to:

- a count function for the occurrence of items with a short-term memory,
- a count function for the occurrence of items with a long-term memory,
- a count function for alternations.

This model includes six free parameters (we use the notations from Kolossa et al.): a decay factor for short-term memory (*β*_S_), two normalized time constants for the dynamic long-term memory (*τ*_1_ and *τ*_2_), the relative weight of item probabilities computed from short-and long-term memory (*α*_S_), the relative weight of probabilities computed for items and their alternations (*α*_∆_), and a parameter capturing the subjective distortion of probabilities (*γ*_∆,2_).

We use the best-fitting values of the free parameters reported by Squires et al. and Kolossa et al. in their respective article.

To compare the fit provided by our models and by the models by Squires et al. and Kolossa et al., we used the Bayesian Information Criterion (BIC). The BIC favors the goodness-of-fit but penalizes model for their number of free parameters [74]. For maximum likelihood estimate of the model parameters and Gaussian residuals, BIC = *n* · log(MSE) + *k* · log(*n*), with *n* the number of fitted data points and *k* the number of free parameters in the model. Note that here, *k* counts the scaling and offset parameter to adjust the model's data and the experimental data, and the internal free parameters of the model (from 0 for fixed belief model with perfect integration, to 6 for the model by Kolossa et al.).

### Summary of the experimental procedure in Huettel et al.

Huettel et al. presented 14 subjects with a stream of two visual stimuli (a square and a circle) randomly generated with equal probability. The sequence length was 1800. Each stimulus was presented for 250 ms, and the stimulus onset asynchrony was 2 s. Subjects were asked to press a dedicated button for each item as quickly as possible. Subjects performed the experiment in an MRI scanner for functional recordings.

### Simulation of the results by Huettel et al

We extracted the data by Huettel et al. from their figure 2 in [2]. We simulated these results using the fixed belief model with leaky integration. We fitted the leak constant to the data using a grid search. We replicated the simulation results with the dynamic belief model, using *p*_c_ = 0.019 for the estimation of frequencies and *p*_c_ = 0.167 for the estimation of transition probabilities, which are the best fitting values for Squires et al. data.

We generated a sequence of 10^5^ stimuli with *p*(X) = 0.5. This large number of observations ensured stable simulation results. Another solution is to generate many short sequences. Both options actually yield similar results because of the limited horizon of the non-stationary estimations used here.

Similar to our fit of Squires et al., we computed posterior inferences and surprise levels. The difference was that surprise levels were sorted based on whether local patterns of stimuli were alternating or repeating, whether the last item violated or continued the pattern, and the length of the pattern (up to 8).

### Summary of the experimental procedure in Cho et al.

Cho et al. presented 6 subjects with a stream of two visual stimuli (a small and a large circle), generated randomly with equal probabilities. Subjects were asked to press a dedicated button for each item as quickly and accurately as possible. Each stimulus was presented until a response was made within a limit of 2 s. The next stimulus appeared after a delay of 0.8 s. Subjects performed 13 series of 120 trials (1560 stimuli in total) with a short break between series.

### Simulation of the results by Cho et al.

We extracted the data by Cho et al. from their figure 1B in [20]. We used the same simulation as for Huettel et al. The only difference being that surprise levels were sorted based on all patterns of alternations and repetitions formed by 5 stimuli.

### Summary of the experimental procedure in Falk et al

As described in [25], Falk presented 219 subjects with sequences of 21 binary visual stimuli (“X” and “O”). The sequences had ratios of alternations ranging from 0.1 to 1 with 0.1 steps. The order of the sequences varied randomly across participants. Each sequence was printed as a row on a paper sheet: stimuli were therefore presented simultaneously. Subjects were asked to rate the apparent randomness of each sequence from 0 to 20, with the indication that this judgment should reflect the likelihood of the sequence having been generated by flipping a fair coin. Ratings were later rescaled between 0 and 1.

### Simulation of the results by Falk

We extracted the data by Falk from figure 1, condition “AR_I_” in [25]. Following the original experiment, we generated sequences of 21 binary stimuli with various probabilities of alternations. For each sequence, we computed the posterior inference of the hidden statistics and the prediction about the next stimulus (the 22_th_) given the previous ones, for each observer model. We used different values for their leak parameter. To quantify the “randomness” of the sequence, we computed the entropy of the prediction: H = – *p*log_2_(*p*) – (1 – *p*)log_2_(1 – *p*). For each leak parameter and alternation frequency, we averaged over 10^4^ sequences to reach stable results. Note that instead of focusing on the last prediction, one could average across successive predictions in each sequence. This alternative yields the same qualitative results as shown in Figure 5.

## Acknowledgements

We thank Tobias Donner, Christophe Pallier and Angela Yu for useful discussions.

This work was funded by INSERM, CEA, Collège de France and by a grant from the European Union Seventh Framework Programme (FP7/2007 2013) under grant agreement no. 604102 (Human Brain Project).

Maxime Maheu is supported by the *Ministère de l’Enseignement Supérieur et de la Recherche* through the *Université Paris Descartes* as well as the *Ecole Doctorale Frontières du Vivant (FdV) – Programme Bettencourt*.

## SUPPLEMENTARY INFORMATION

***Supplementary Figure 1: Goodness-of-fit of electrophysiological data for different parameter values and models***

The plot shows the mean squared error of model fit, for different models, different values of their free parameter and different datasets. The inset shows a zoom around the best-fitting parameter. Note that different models have different inference styles: fixed belief with perfect integration within a window of observation, fixed belief with leaky integration and dynamic belief (presented in different columns) and they estimate different statistics: item frequency, alternation frequency and transition probabilities (presented as colored lines within each plot).

***Supplementary Figure 2: Sequential effects in reaction times predicted by the dynamic belief model***

This figure is similar to Figure 4. The only difference is that theoretical surprise levels are computed from the dynamic belief model (see Methods). The free parameter of the models, the *a priori* change probability for the estimated statistics, was selected independently, as the best fitting value for Squires et al. data.

***Supplementary Figure 3: Judgement of randomness predicted by the dynamic belief model***

This figure is similar to Figure 5. The only difference is that theoretical entropy levels are computed from the dynamic belief model (see Methods) using different a priori change probabilities for the estimated statistics.

***Supplementary Figure 4: The models make qualitatively different predictions***

We estimated whether the models we consider make predictions that can be distinguished quantitatively from one another. We started by simulating the results of the Squires et al experiment, using the best-fitting parameters of each model. As in Squires et al, each simulated data set contained 48 data points, corresponding to 16 patterns times 3 block types. We then estimated the ability of a given model to recover the predictions of another model with a leave-one out procedure. More precisely, we took the 48 simulated values of a given model and we adjusted the free parameters of another model to 47 of these simulated values, leaving one out. Given these fitted parameters, we then computed the prediction of the second model about the left-out point and we measured the error (the unsigned difference) compared with the original simulation. The bars show, for all pairs of models, the mean error and SEM across left-out points. The comparison of a model against itself yields no error (see arrows). Different models that make similar predictions should yield an error close to 0. This occurs in only one case (see dashed arrows): the predictions of a model learning perfectly a given statistic can be recovered almost exactly by a model learning the same statistic with a leaky integration. This is because the leak can be adjusted so as to approach a perfect integration. Critically, models that learn different statistics all produced quantitatively different predictions.

***Supplementary Figure 5:* Comparison of the predictive accuracy of different models (Squires et al dataset).**

We estimated the predictive accuracy of each model using a leave-one out procedure. The parameters of each model (the offset and slope of the linear transformation from theoretical surprise to P300 data, and the leak of leaky integration models) were fitted to all data points but one. We then measured the error of a given model, as the distance between the actual left-out point and the value predicted by the model and its fitted parameters. Since Squires et al report data for 16 patterns in 3 different block types, we repeated the leave-one out procedure 16*3=48 times. Bars show the mean error with SEM, and the numbers indicate the mean (and SEM) of the best-fitting leak parameter ω across left-out points. The model learning local estimates of transition probabilities achieved the best predictive accuracy, i.e. the smallest error.

## Supplementary equations

Several statistics are embedded in the space of transition probabilities. Indeed, transition probabilities fully specify the frequency of items:

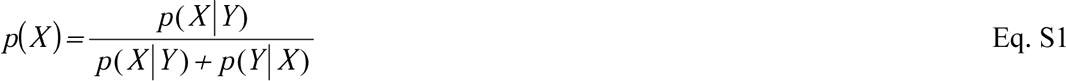

They also fully specify the frequency of ordered pairs of items:

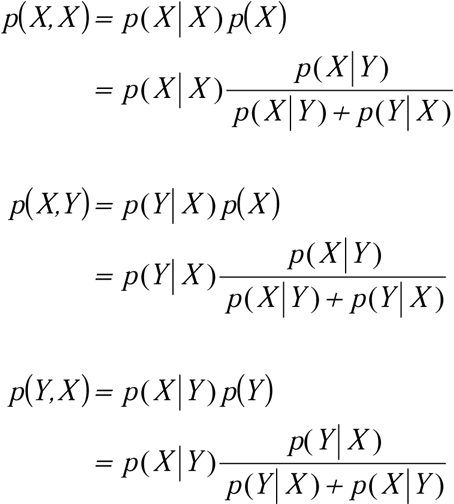

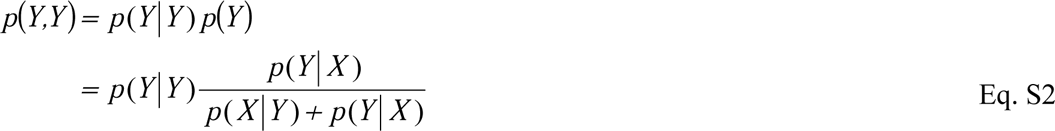

And they fully specify the frequency of alternations:

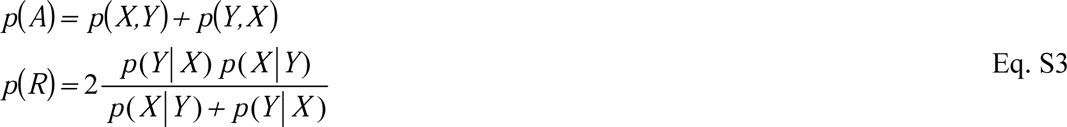

## References

1. Squires KC, Wickens C, Squires NK, Donchin E. The effect of stimulus sequence on the waveform of the cortical event-related potential. Science. 1976;193: 1142–1146. doi:10.1126/science.959831

2. Huettel SA, Mack PB, McCarthy G. Perceiving patterns in random series: dynamic processing of sequence in prefrontal cortex. Nat Neurosci. 2002;5: 485–490. doi:10.1038/nn841

3. Bendixen A, Roeber U, Schröger E. Regularity Extraction and Application in Dynamic Auditory Stimulus Sequences. J Cogn Neurosci. 2007;19: 1664–1677. doi:10.1162/jocn.2007.19.10.1664

4. Mars RB, Debener S, Gladwin TE, Harrison LM, Haggard P, Rothwell JC, et al. Trial-by-Trial Fluctuations in the Event-Related Electroencephalogram Reflect Dynamic Changes in the Degree of Surprise. J Neurosci. 2008;28: 12539–12545. doi:10.1523/JNEUROSCI.2925-08.2008

5. Bekinschtein TA, Dehaene S, Rohaut B, Tadel F, Cohen L, Naccache L. Neural signature of the conscious processing of auditory regularities. Proc Natl Acad Sci. 2009;106: 1672–1677. doi:10.1073/pnas.0809667106

6. Kimura M, Schröger E, Czigler I, Ohira H. Human visual system automatically encodes sequential regularities of discrete events. J Cogn Neurosci. 2010;22: 1124–1139.

7. Wacongne C, Changeux J-P, Dehaene S. A Neuronal Model of Predictive Coding Accounting for the Mismatch Negativity. J Neurosci. 2012;32: 3665–3678. doi:10.1523/JNEUROSCI.5003-11.2012

8. Yaron A, Hershenhoren I, Nelken I. Sensitivity to Complex Statistical Regularities in Rat Auditory Cortex. Neuron. 2012;76: 603–615. doi:10.1016/j.neuron.2012.08.025

9. Kolossa A, Fingscheidt T, Wessel K, Kopp B. A Model-Based Approach to Trial-By-Trial P300 Amplitude Fluctuations. Front Hum Neurosci. 2013;6. doi:10.3389/fnhum.2012.00359

10. Lieder F, Daunizeau J, Garrido MI, Friston KJ, Stephan KE. Modelling Trial-by-Trial Changes in the Mismatch Negativity. Sporns O, editor. PLoS Comput Biol. 2013;9: e1002911. doi:10.1371/journal.pcbi.1002911

11. Strauss M, Sitt JD, King J-R, Elbaz M, Azizi L, Buiatti M, et al. Disruption of hierarchical predictive coding during sleep. Proc Natl Acad Sci U S A. 2015;112: E1353–1362. doi:10.1073/pnas.1501026112

12. Hyman R. Stimulus information as a determinant of reaction time. J Exp Psychol. 1953;45: 188–196. doi:10.1037/h0056940

13. Bertelson P. Sequential redundancy and speed in a serial two-choice responding task. Q J Exp Psychol. 1961;13: 90–102. doi:10.1080/17470216108416478

14. Tune SG. Response preferences: A review of some relevant literature. Psychol Bull. 1964;61: 286–302. doi:10.1037/h0048618

15. Rouanet H. Les modèles stochastiques d’apprentissage, Recherches et perspectives [Internet]. Paris: Mouton; 1967. Available: http://www.degruyter.com/view/product/150496

16. Schvaneveldt RW, Chase WG. Sequential effects in choice reaction time. J Exp Psychol. 1969;80: 1–8. doi:10.1037/h0027144

17. Kirby NH. Sequential effects in two-choice reaction time: automatic facilitation or subjective expectancy? J Exp Psychol Hum Percept Perform. 1976;2: 567–577.

18. Soetens E, C L, E J. Expectancy or automatic facilitation? Separating sequential effects in two-choice reaction time. J Exp Psychol Hum Percept Perform. 1985;11: 598–616. doi:10.1037/0096-1523.11.5.598

19. Sommer W, Leuthold H, Soetens E. Covert signs of expectancy in serial reaction time tasks revealed by event-related potentials. Percept Psychophys. 1999;61: 342–353. doi:10.3758/BF03206892

20. Cho RY, Nystrom LE, Brown ET, Jones AD, Braver TS, Holmes PJ, et al. Mechanisms underlying dependencies of performance on stimulus history in a two-alternative forced-choice task. Cogn Affect Behav Neurosci. 2002;2: 283–299. doi:10.3758/CABN.2.4.283

21. Lungu OV, Wächter T, Liu T, Willingham DT, Ashe J. Probability detection mechanisms and motor learning. Exp Brain Res. 2004;159: 135–150. doi:10.1007/s00221-004-1945-7

22. Perruchet P, Cleeremans A, Destrebecqz A. Dissociating the effects of automatic activation and explicit expectancy on reaction times in a simple associative learning task. J Exp Psychol Learn Mem Cogn. 2006;32: 955–965. doi:10.1037/0278-7393.32.5.955

23. Yu AJ, Cohen JD. Sequential effects: Superstition or rational behavior? Adv Neural Inf Process Syst. 2008;21: 1873–1880.

24. Kareev Y. Positive bias in the perception of covariation. Psychol Rev. 1995;102: 490–502. doi:10.1037/0033-295X.102.3.490

25. Falk R, Konold C. Making sense of randomness: Implicit encoding as a basis for judgment. Psychol Rev. 1997;104: 301.

26. Hahn U, Warren PA. Perceptions of randomness: Why three heads are better than four. Psychol Rev. 2009;116: 454–461. doi:10.1037/a0015241

27. Sun Y, Wang H. Perception of randomness: On the time of streaks. Cognit Psychol. 2010;61: 333–342. doi:10.1016/j.cogpsych.2010.07.001

28. Fawcett TW, Fallenstein B, Higginson AD, Houston AI, Mallpress DEW, Trimmer PC, et al. The evolution of decision rules in complex environments. Trends Cogn Sci. 2014;18: 153–161. doi:10.1016/j.tics.2013.12.012

29. Sun Y, O’Reilly RC, Bhattacharyya R, Smith JW, Liu X, Wang H. Latent structure in random sequences drives neural learning toward a rational bias. Proc Natl Acad Sci. 2015;112: 3788–3792. doi:10.1073/pnas.1422036112

30. Deneve S, Latham PE, Pouget A. Reading population codes: a neural implementation of ideal observers. Nat Neurosci. 1999;2: 740–745.

31. Rao RP. An optimal estimation approach to visual perception and learning. Vision Res. 1999;39: 1963–1989.

32. Ernst MO, Banks MS. Humans integrate visual and haptic information in a statistically optimal fashion. Nature. 2002;415: 429–433. doi:10.1038/415429a

33. Beck JM, Ma WJ, Kiani R, Hanks T, Churchland AK, Roitman J, et al. Probabilistic Population Codes for Bayesian Decision Making. Neuron. 2008;60: 1142–1152. doi:10.1016/j.neuron.2008.09.021

34. Maloney LT, Zhang H. Decision-theoretic models of visual perception and action. Vision Res. 2010;50: 2362–2374. doi:10.1016/j.visres.2010.09.031

35. Berkes P, Orbán G, Lengyel M, Fiser J. Spontaneous cortical activity reveals hallmarks of an optimal internal model of the environment. Science. 2011;331: 83–87. doi:10.1126/science.1195870

36. Girshick AR, Landy MS, Simoncelli EP. Cardinal rules: visual orientation perception reflects knowledge of environmental statistics. Nat Neurosci. 2011;14: 926–932. doi:10.1038/nn.2831

37. Diaconescu AO, Mathys C, Weber LAE, Daunizeau J, Kasper L, Lomakina EI, et al. Inferring on the intentions of others by hierarchical Bayesian learning. PLoS Comput Biol. 2014;10: e1003810. doi:10.1371/journal.pcbi.1003810

38. Hoyer PO, Hyvärinen A. Interpreting Neural Response Variability as Monte Carlo Sampling of the Posterior. Advances in Neural Information Processing Systems. MIT Press; 2002. p. 2002.

39. Lee TS, Mumford D. Hierarchical Bayesian inference in the visual cortex. J Opt Soc Am A Opt Image Sci Vis. 2003;20: 1434–1448.

40. Knill DC, Pouget A. The Bayesian brain: the role of uncertainty in neural coding and computation. Trends Neurosci. 2004;27: 712–719. doi:10.1016/j.tins.2004.10.007

41. Ma WJ, Beck JM, Latham PE, Pouget A. Bayesian inference with probabilistic population codes. Nat Neurosci. 2006;9: 1432–1438. doi:10.1038/nn1790

42. Fiser J, Berkes P, Orbán G, Lengyel M. Statistically optimal perception and learning: from behavior to neural representations. Trends Cogn Sci. 2010;14: 119–130. doi:10.1016/j.tics.2010.01.003

43. Ma WJ, Jazayeri M. Neural coding of uncertainty and probability. Annu Rev Neurosci. 2014;37: 205–220. doi:10.1146/annurev-neuro-071013-014017

44. Jaynes ET. Probability Theory: The Logic of Science. Cambridge University Press; 2003.

45. Friston K. Learning and inference in the brain. Neural Netw Off J Int Neural Netw Soc. 2003;16: 1325–1352. doi:10.1016/j.neunet.2003.06.005

46. Friston K. The free-energy principle: a unified brain theory? Nat Rev Neurosci. 2010;11: 127–138. doi:10.1038/nrn2787

47. Dehaene S, Meyniel F, Wacongne C, Wang L, Pallier C. The Neural Representation of Sequences: From Transition Probabilities to Algebraic Patterns and Linguistic Trees. Neuron. 2015;88: 2–19. doi:10.1016/j.neuron.2015.09.019

48. Wilder M, Jones M, Mozer MC. Sequential effects reflect parallel learning of multiple environmental regularities. In: Bengio Y, Schuurmans D, Lafferty JD, Williams CKI, Culotta A, editors. Advances in Neural Information Processing Systems 22. Curran Associates, Inc.; 2009. pp. 2053–2061. Available: http://papers.nips.cc/paper/3870-sequential-effects-reflect-parallel-learning-of-multiple-environmental-regularities.pdf

49. Shannon CE. A mathematical theory of communication. Bell Syst Tech J. 1948;27: 379–423.

50. Gelman A, Carlin JB, Stern HS, Dunson DB, Vehtari A, Rubin DB. Bayesian Data Analysis, Third Edition. CRC Press; 2013.

51. Rao RP, Sejnowski TJ. Predictive coding, cortical feedback, and spike-timing dependent plasticity. Statistical Theories of the Brain. 2000.

52. Friston K. A theory of cortical responses. Philos Trans R Soc Lond B Biol Sci. 2005;360: 815–836. doi:10.1098/rstb.2005.1622

53. Wolpert DM, Ghahramani Z. Computational principles of movement neuroscience. Nat Neurosci. 2000;3: 1212–1217. doi:10.1038/81497

54. Todorov E. Optimality principles in sensorimotor control. Nat Neurosci. 2004;7: 907–915. doi:10.1038/nn1309

55. Sutton RS, Barto AG. Introduction to Reinforcement Learning. 1st ed. Cambridge, MA, USA: MIT Press; 1998.

56. Behrens TEJ, Woolrich MW, Walton ME, Rushworth MFS. Learning the value of information in an uncertain world. Nat Neurosci. 2007;10: 1214–1221. doi:10.1038/nn1954

57. Nassar MR, Rumsey KM, Wilson RC, Parikh K, Heasly B, Gold JI. Rational regulation of learning dynamics by pupil-linked arousal systems. Nat Neurosci. 2012;15: 1040–1046. doi:10.1038/nn.3130

58. Ossmy O, Moran R, Pfeffer T, Tsetsos K, Usher M, Donner TH. The Timescale of Perceptual Evidence Integration Can Be Adapted to the Environment. Curr Biol. 2013;23: 981–986. doi:10.1016/j.cub.2013.04.039

59. McGuire JT, Nassar MR, Gold JI, Kable JW. Functionally Dissociable Influences on Learning Rate in a Dynamic Environment. Neuron. 2014;84: 870–881. doi:10.1016/j.neuron.2014.10.013

60. Meyniel F, Schlunegger D, Dehaene S. The Sense of Confidence during Probabilistic Learning: A Normative Account. PLoS Comput Biol. 2015;11: e1004305. doi:10.1371/journal.pcbi.1004305

61. Wang X-J. Probabilistic Decision Making by Slow Reverberation in Cortical Circuits. Neuron. 2002;36: 955–968. doi:10.1016/S0896-6273(02)01092-9

62. Honey CJ, Thesen T, Donner TH, Silbert LJ, Carlson CE, Devinsky O, et al. Slow Cortical Dynamics and the Accumulation of Information over Long Timescales. Neuron. 2012;76: 423–434. doi:10.1016/j.neuron.2012.08.011

63. Gallistel CR, Krishan M, Liu Y, Miller R, Latham PE. The perception of probability. Psychol Rev. 2014;121: 96–123. doi:10.1037/a0035232

64. Kemp C, Tenenbaum JB. The discovery of structural form. Proc Natl Acad Sci U S A. 2008;105: E10687–10692. doi:10.1073/pnas.0802631105

65. Saffran JR, Aslin RN, Newport EL. Statistical learning by 8-month-old infants. Science. 1996;274: 1926–1928.

66. Meyer T, Ramachandran S, Olson CR. Statistical Learning of Serial Visual Transitions by Neurons in Monkey Inferotemporal Cortex. J Neurosci. 2014;34: 9332–9337. doi:10.1523/JNEUROSCI.1215-14.2014

67. Ramachandran S, Meyer T, Olson CR. Prediction suppression in monkey inferotemporal cortex depends on the conditional probability between images. J Neurophysiol. 2016;115: 355–362. doi:10.1152/jn.00091.2015

68. Summerfield C, Trittschuh EH, Monti JM, Mesulam M-M, Egner T. Neural repetition suppression reflects fulfilled perceptual expectations. Nat Neurosci. 2008;11: 1004–1006. doi:10.1038/nn.2163

69. Todorovic A, van Ede F, Maris E, de Lange FP. Prior Expectation Mediates Neural Adaptation to Repeated Sounds in the Auditory Cortex: An MEG Study. J Neurosci. 2011;31: 9118–9123. doi:10.1523/JNEUROSCI.1425-11.2011

70. Bornstein AM, Daw ND. Cortical and Hippocampal Correlates of Deliberation during Model-Based Decisions for Rewards in Humans. PLoS Comput Biol. 2013;9: e1003387. doi:10.1371/journal.pcbi.1003387

71. Rose M, Haider H, Büchel C. Unconscious detection of implicit expectancies. J Cogn Neurosci. 2005;17: 918–927.

72. van Zuijen TL, Simoens VL, Paavilainen P, Näätänen R, Tervaniemi M. Implicit, Intuitive, and Explicit Knowledge of Abstract Regularities in a Sound Sequence: An Event-related Brain Potential Study. J Cogn Neurosci. 2006;18: 1292–1303. doi:10.1162/jocn.2006.18.8.1292

73. Atas A, Faivre N, Timmermans B, Cleeremans A, Kouider S. Nonconscious Learning From Crowded Sequences. Psychol Sci. 2014;25: 113–119. doi:10.1177/0956797613499591

74. Schwarz G. Estimating the Dimension of a Model. Ann Stat. 1978;6: 461–464. doi:10.1214/aos/1176344136

